# Exposure to spirochete-infected *Ornithodoros turicata* collected from the home of an individual in the Austin, Texas Metropolitan Area

**DOI:** 10.1101/2025.11.12.688062

**Authors:** Serhii Filatov, Bonny Mayes, Sarah M. Gunter, Alexander R. Kneubehl, Job E. Lopez

**Author notes:** **Corresponding author:** Job E. Lopez. Contributed equally to this manuscript (Serhii Filatov, Bonny Mayes).

## Abstract

A Central Texas resident found an engorging *Ornithodoros turicata* tick attached to their leg while at home. The tick was submitted for testing, which confirmed that it was infected with a *Borrelia* species. Spirochetes were successfully cultured and whole genome sequencing confirmed that the species was *Borrelia turicatae*. The tick submitter reported being hospitalized and was recovering from Bell’s Palsy. Searching the Texas National Electronic Disease Surveillance System indicated that the individual had an undiagnosed febrile illness that spanned three weeks prior to finding the tick attached on them. These findings indicated possible recurrent tick exposure in a domestic environment.

## Introduction

In the United States, soft tick relapsing fever (STRF) spirochetes predominantly circulate among wild vertebrates and cavity-dwelling argasid tick vectors. Individuals that enter the natural habitat of *Ornithodoros* spp. and are bitten typically become symptomatic within three to seven days ^1^. Due to the rapid feeding behavior of the ticks, they are seldom found attached to their hosts, and exposed individuals are usually unaware of being bitten.

The predominant cause of STRF in the southwestern United States is *Borrelia turicatae*, transmitted by *Ornithodoros turicata* ^2^. Historically, human cases of STRF have been linked to caves and rural areas like cabins and outbuildings frequented by peridomestic mammals ^3^. Recent reports indicate a shift in vector ecology, with spirochete-infected *O. turicata* identified in public parks and green spaces within urban centers in Texas ^4, 5^. In this current study, we report an individual discovering an attached spirochete-infected *O. turicata* while in their home, underscoring the potential for intra-household transmission.

## Methods

In early March 2024, the Texas Department of State Health Services (DSHS) Zoonosis Control Branch (ZCB) received a tick from an Austin Metropolitan area resident for testing at University of North Texas Health Fort Worth (UNT Health). The ZCB has been offering free tick testing through UNT Health for Texas residents since 2004. The submitter reported that the tick was attached to their ankle when they sat down on their couch. Upon examination, ZCB staff noted that the tick was an engorged, live argasid (soft) tick, likely *O. turicata* and a vector of STRF. The tick was sent to Baylor College of Medicine (BCM) since UNT Health does not test for *B. turicatae*.

At BCM, the tick was examined morphologically and for infectivity. Using a stereomicroscope, the ventral surface was visualized to identify developmental features. To determine infectivity of the tick, animal studies were performed. All animal work was approved through BCM’s Institutional Animal Care and Use Committee. The tick was fed on a 6-week-old Institute of Cancer Research (ICR) mouse, and a drop of blood was collected from the animal daily and inspected for spirochetes by dark field microscopy. When spirochetes were visualized, the animal was sedated, exsanguinated, and the collected blood was centrifuged at 500 × g. The spirochete-enriched serum was inoculated into 4 mL of modified Barbour-Stonner-Kelly (mBSK) medium ^6^. Once spirochetes reached mid-exponential growth, 40 ml of fresh mBSK medium was inoculated, and glycerol stocks were made. The 40 ml culture was centrifuged at 11,000 x g for 20 min at 4 °C to pellet spirochetes, and total DNA was extracted using the phenol-chloroform method ^7^. Using an established approach of Illumina and Oxford Nanopore Technologies ^8^, the isolate genome was sequenced and assembled for phylogenomic analysis.

## Results

### Identification and characterization of the tick attached to a human

Physical examination determined the tick’s developmental stage. When the tick arrived at BCM from DSHS it was still engorged. The absence of a genital aperture, a structure that develops during the transition from nymph to adult, indicated that the specimen was an immature tick. Over the following three weeks, the tick molted. A follow-up morphological examination identified the presence of a genital aperture characteristic of an adult male *Ornithodoros* tick (Supplementary Figure S1). These findings indicated that the individual was fed upon by a nymphal *O. turicata* that molted into an adult male.

### Evaluation of the infection status of the O. turicata tick

After feeding the tick on a clean laboratory mouse, spirochetes were visualized in the blood four days later. The spirochetes were successfully cultured in mBSK and the isolate was designated ZLK. The ZLK isolate could be resurrected from glycerol stocks and grown after cryopreservation.

### Genomic assessment of the spirochete isolate

Using short- and long-read sequencing technology, the genome of ZLK was generated. The merqury tool (v1.3) determined the assembly was 99.7% complete and had a consensus quality of +inf, indicating that errors were not detected in the assembly. The genome assembly was 1,670,619 base pairs in length and consisted of the linear chromosome, 12 linear plasmids, and six circular plasmids. A phylogenomic analysis confirmed that the ZLK isolate was *B. turicatae* (Figure 1 A). Plasmids families were typed using the genes that encode the PF32 and PF57/62 proteins (Figure 1 B). ZLK grouped most closely with a human (BTE5EL) and tick (91E135) isolate from Texas, and a tick isolate from Mexico (BTCAM1) ^8, 9^.

**Figure 1.**
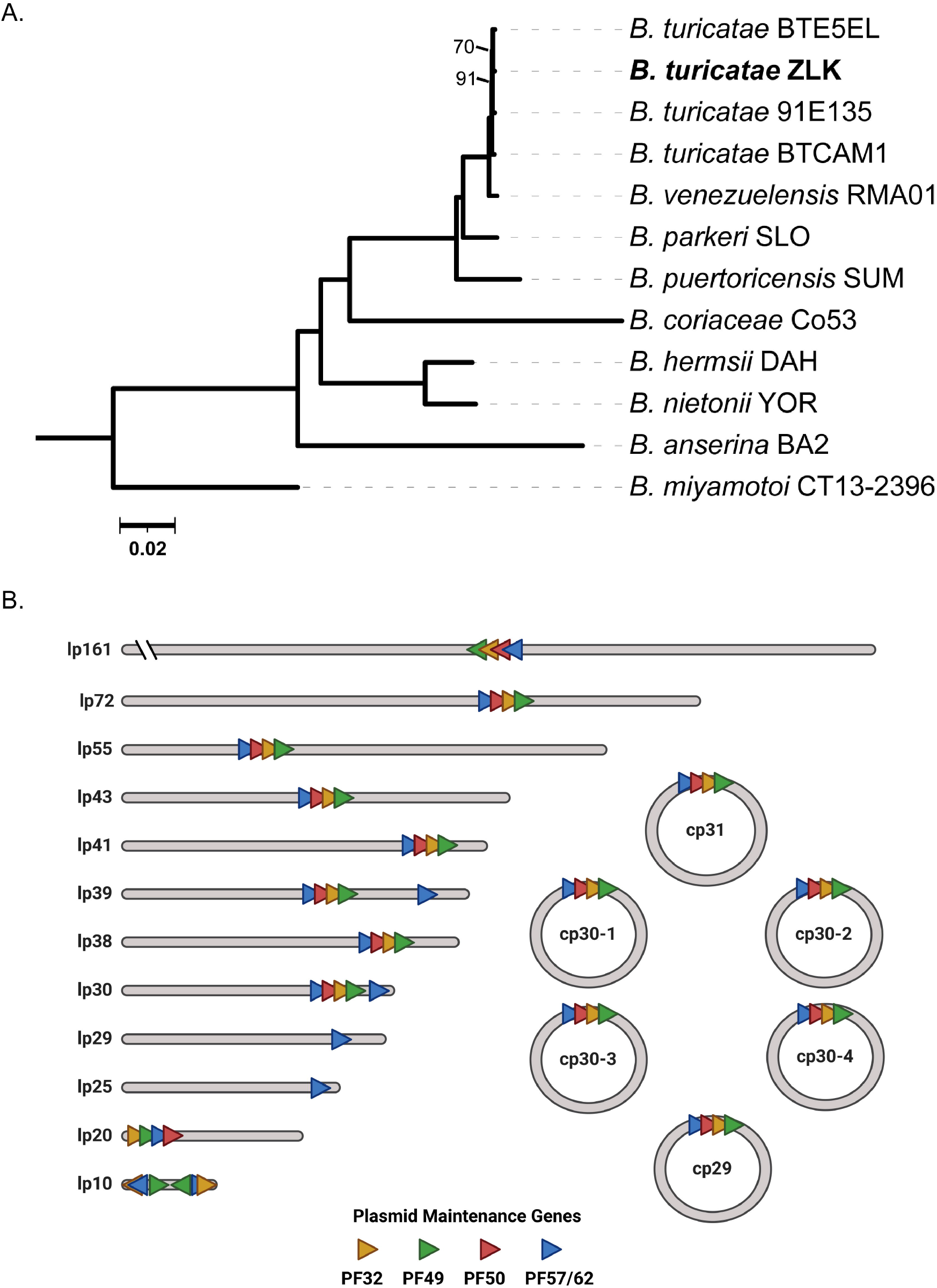
Phylogenomic (A) and plasmid analysis (B) of the ZLK isolate. The maximum-likelihood tree generated with an edge-linked proportional partition model (1,000 ultra-fast bootstrap replicates) grouped the ZLK isolate (boldface) with other *B. turicatae* strains. Scale bar indicates 0.02 substitutions per site. Assembly of the genome identified 12 linear plasmids and six circular plasmids. The plasmid partitioning genes thought to be responsible for the heritable maintenance of the plasmids are indicated in colored triangles with gene orientation indicated by the direction of the triangle’s point.

### Clinical history of the tick submitter

After the tick was identified at DSHS, ZCB called the submitter to inform them that the tick was a potential vector of STRF spirochetes. The individual noted that prior to finding the attached tick, they were recently hospitalized and still recovering from Bell’s palsy (Table 1). Additionally, they reported testing positive for Lyme disease. Since individuals infected with STRF spirochetes can test positive for Lyme disease and both diseases are reportable in Texas, medical records were requested. In addition, ZCB searched Texas National Electronic Disease Surveillance System (NEDSS) for electronic lab reports (ELRs) on the submitter. Medical records revealed that the submitter tested positive on the first tier of the Lyme disease serology test (ELISA), while the second tier (immunoblots) was negative. Serology for flea-borne typhus (day 5 and 11 serum samples) and spotted fever rickettsiosis (day 5 serum sample) were negative; influenza, SARS-CoV-2, strep throat, West Nile virus, syphilis, and HIV test results were also negative. A day 11 serum sample was obtained and forwarded to CDC for STRF Western blot testing. The sample was positive for recombinant glycerophosphodiester phosphodiesterase (rGlpQ), an antigen that allows for serologic differentiation between relapsing fever and Lyme borreliosis ^10^. However, antibodies to a 17kDa diagnostic antigen were undetectable. CDC noted that false negative results may occur from serum samples collected <14 days after symptom onset. Unfortunately, attempts to re-contact and obtain another serum sample from the individual were unsuccessful.

## Discussion

The clinical presentation and absence of an alternate explanation for the tick submitter’s illness suggests infection with STRF spirochetes. However, the individual’s location of exposure to *O. turicata* prior to illness remains unclear. They did not report travel outside of Texas in the previous three months, yet frequented Austin city parks. Exposure may have occurred at a park, but they denied finding an attached tick on their body prior to the onset of disease. Evidence for intrahousehold exposure included the discovery of an engorging *O. turicata* tick 17 days after their symptoms began (Table 1). If this tick was responsible for the individual’s original illness, there was sufficient time for the tick to molt and subsequently feed again 17 days later. However, it remains unclear how the tick was introduced into the individual’s dwelling or if there was an underlying infestation of *O. turicata*.

A limitation of the serological analysis is that the identity of 17 kDa diagnostic antigen and the antibody kinetics against this protein during natural infection are unknown. Consequently, it is unclear whether IgG responses would have been detectable when the patient’s serum sample was collected 11 days post symptom onset. In contrast, the detection of anti-rGlpQ IgG antibodies is noteworthy. GlpQ is highly antigenic ^11^, and it was evident that anti-GlpQ IgG antibodies were circulating when the patient’s serum sample was collected. When serological results are questionable, microscopy, PCR, and bacterial culture would have supported a more conclusive diagnosis.

Our findings contribute to growing evidence that STRF spirochetes are endemic in urban areas in Travis County and highlight the need for increased awareness among clinicians. Over the past seven years *B. turicatae* infected ticks have been collected and associated with human exposure in densely population regions of Austin, Texas (Supplementary Figure S2) ^4, 5, 12, 13^. With greater ecological overlap between humans, *O. turicata*, and vertebrates, ticks are likely to enter domestic environments and feed on humans, as previously observed ^14^. Once introduced, an endemic focus of infected *O. turicata* can persist for decades because these ticks live over 10 years, can feed over 15 times throughout their lifespan, and vertically transmit STRF spirochetes ^15^. There is ample opportunity for *O. turicata* to acquire and transmit *B. turicatae*. Consequently, we recommend increased surveillance efforts in green spaces, neighborhoods, and parks to identify *O. turicata* and the vertebrates that aid in maintaining *B. turicatae*.

## Acknowledgments and funding

Amber Frenzel with DSHS Region 7 Zoonosis Control for obtaining medical records. This project was funded in part by Lyda Hill Philanthropies, Baylor College of Medicine Interim Funding Program, and the Texas EcoLab Program.

## Notes

### Competing Interest Statement

The authors have declared no competing interest.

