## Supplementary Figures for "Exposure to spirochete-infected *Ornithodoros turicata* collected from the home of an individual in the Austin, Texas Metropolitan Area"

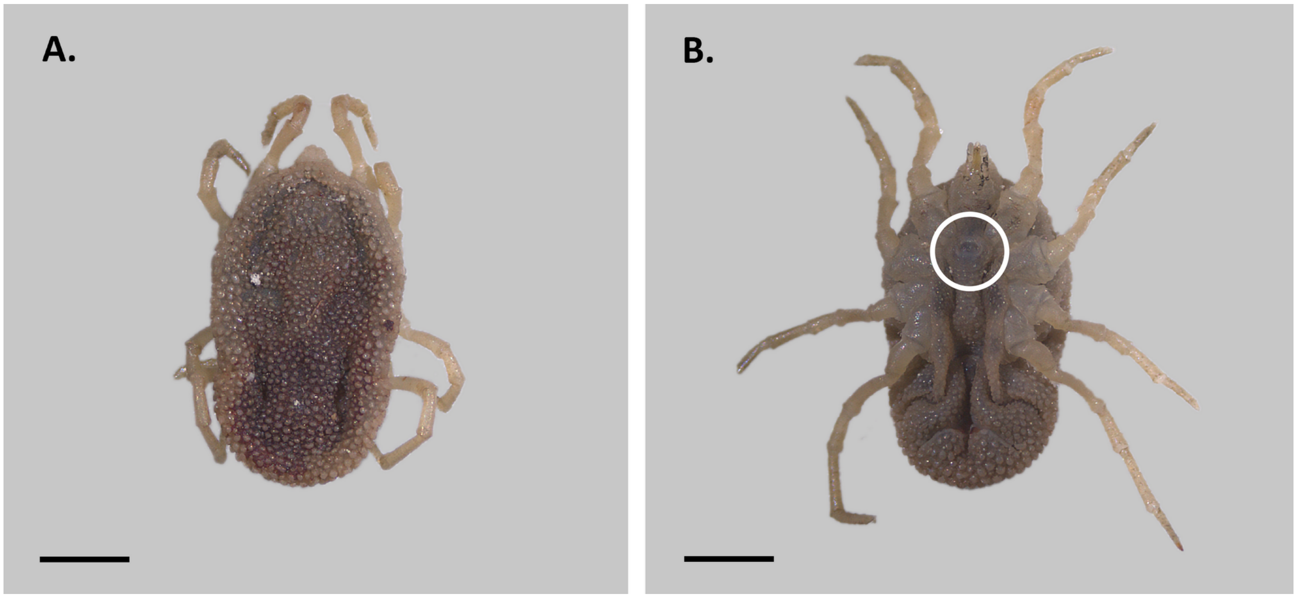


Supplementary Figure S1. Morphological characterization of *O. turicata*. Shown are the dorsal (A) and ventral (B) surface of the tick. The genital aperture is shown within the white circle (B). The scale bar represents 1 mm.

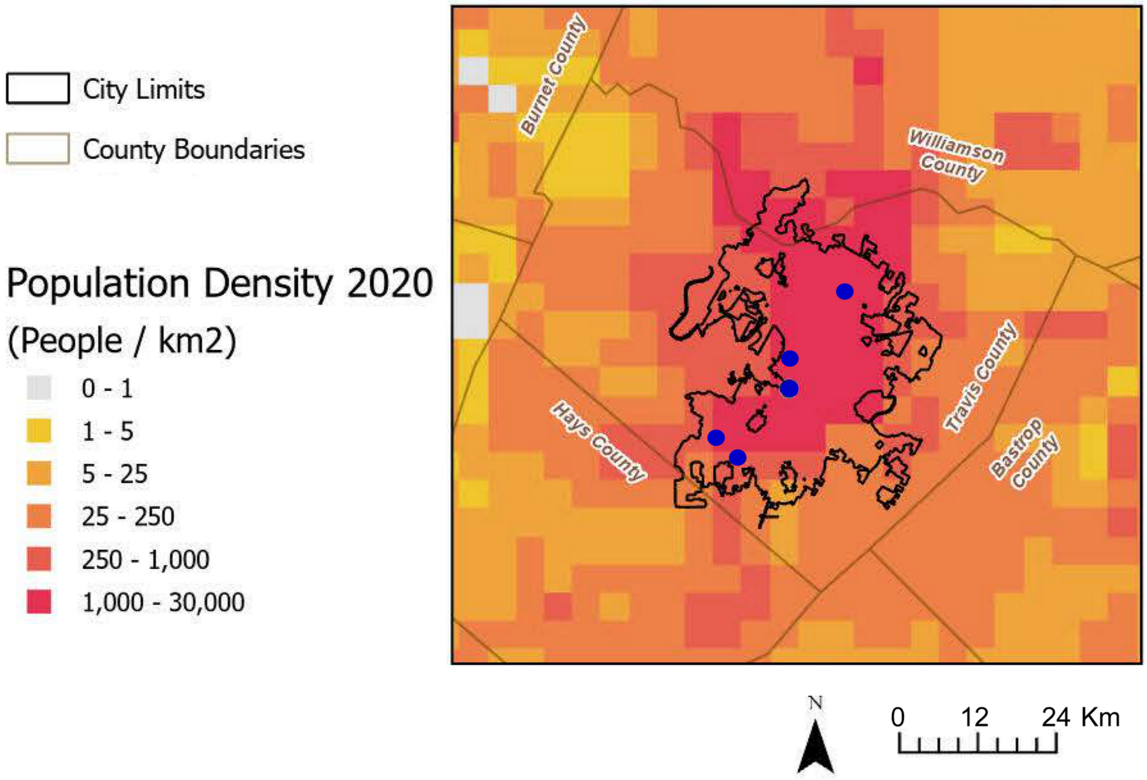


Supplementary Figure S2. Population density map of the Austin, Texas region. The city limits (black outline) and county boundaries (grey outline) are shown. Each blue dot represents collection sites of infected *O. turicata* ticks. The map was generated in ArcGISPro version 3.2.
